# Rapid adaptation often occurs through mutations to the most highly conserved positions of the RNA polymerase core enzyme

**DOI:** 10.1101/2021.11.16.468808

**Authors:** Yasmin Cohen, Ruth Hershberg

## Abstract

Mutations to the genes encoding the RNA polymerase core enzyme (RNAPC) and additional housekeeping regulatory genes were found to be involved in rapid adaptation, in the context of numerous evolutionary experiments, in which bacteria were exposed to diverse selective pressures. This provides a conundrum, as the housekeeping genes that were so often mutated in response to these diverse selective pressures tend to be among the genes that are most conserved in their sequences across the bacterial phylogeny. In order to further examine this apparent discrepancy, we characterized the precise positions of the RNAPC involved in adaptation to a large variety of selective pressures. We found that different positions of the RNAPC are involved in adaptation to various stresses, with very little overlap found between stresses. We further found that RNAPC positions involved in adaptation tended to be more evolutionary conserved, were more likely to occur within defined protein domains, and tended to be closer to the complex’s active site, compared to all other RNAPC positions. Finally, we could show that this observed trend of higher conservation of positions involved in rapid adaptation extends beyond the RNAPC to additional housekeeping genes. Combined, our results demonstrate that the positions that change most readily in response to well defined selective pressures exerted in lab environments are also those that evolve most slowly in nature. This suggests that such adaptations may not readily occur in nature, due to their antagonistically pleiotropic effects, or that if they do occur in nature, they are highly transient.

## Introduction

Evolutionary experiments have been instrumental in enabling researchers to study evolution as it happens within controlled environments, and particularly in enabling the study of bacterial rapid adaptation (Kawecki, et al. 2012; Barrick and Lenski 2013; Katz, et al. 2021). Bacteria in particular are useful for evolutionary experiments, because they have short generation times, enabling to study their evolution over relatively large numbers of generations, in a relatively short amount of time. Many bacterial species can be frozen and later revived, enabling researchers to go “back in time” and compare an evolved strain to its ancestor. During evolutionary experiments bacterial populations are exposed to specific selective pressures and the manner in which they adapt to these pressures is examined. The advent of next generation whole genome sequencing technologies enabled many studies that characterized the adaptive mutations that occur in response to specific selective pressures. Given that *Escherichia coli* is the most commonly used bacterial model organisms, a substantial fraction of such studies were carried out in *E. coli*.

Evolutionary experiments have highlighted the remarkable capability of bacteria to undergo rapid adaptation. Such rapid adaptation often occurs through mutations to very central housekeeping genes (reviewed in (Hershberg 2017) and (Maddamsetti, et al. 2017)). The most obvious example of this trend is adaptations occurring within the RNA polymerase core enzyme (RNAPC) genes, *rpoB* and *rpoC*. The *rpoB* gene encodes the RNAPC’s β subunit and *rpoC* encodes its β’ subunit. These two subunits occupy 80% of the total mass of the core enzyme and together form its active site (Sutherland and Murakami 2018). Mutations within *rpoB* and *rpoC* were shown to be involved in adaptation to a variety of selective pressures including exposure to lethal doses of antibiotics (Severinov, et al. 1993; Reynolds 2000; Delgado, et al. 2001; Srivastava, et al. 2012; Degen, et al. 2014), high temperatures (Tenaillon, et al. 2012), low nutrients (Conrad, et al. 2010), exposure to radiation (Bruckbauer, et al. 2019), and prolonged resource exhaustion (Avrani, et al. 2017; Gross, et al. 2020).

The fact that housekeeping genes such as the RNAPC tend to rapidly acquire adaptive mutations, altering their sequences, in response to a large variety of selective pressures, stands in apparent contrast to the high levels of conservation of these genes. Housekeeping genes in general and the RNAPC in particular tend to be extremely well conserved in their sequences, structure and function from bacteria to humans (Archambault and Friesen 1993; Zhang, et al. 1999). This conservation is extensive enough to allow the bacterial RNA polymerase to serve as a model for understanding the basic principles at work in all cellular RNA polymerases (Borukhov and Nudler 2008). Within bacteria the sequences of *rpoB* and *rpoC* are conserved enough to enable their usage as a slowly evolving gene markers in the study of bacterial phylogeny (Lan, et al. 2016).

Here, we characterize the positions of the RNAPC involved in known adaptations and compare them to positions in which no such adaptations have so far been observed. This allows us to show that positions involved in adaptation tend to be even more conserved than other positions of the RNAPC and tend to be located more closely to the protein complex’s active site. The finding that adaptations tend to occur within more conserved positions also extends to additional proteins. Our results further indicate that a unique set of RNAPC positions are involved in adaptation to different conditions. While adaptations in general tend to occur close to the RNAPC active site, adaptations to different conditions tend to cluster to different parts of the complex. Outliers to these reported trends are adaptations occurring in response to heat shock, which do not tend to occur within more conserved positions, and which do not tend to cluster close to each other or to the protein’s active site.

## Materials and Methods

### Datasets

*E*.*coli* K12 MG1655 protein sequences were downloaded from the National Center for Biotechnology Information (NCBI) database (May 2019) (Coordinators 2018). The protein sequences of 44,048 fully sequenced bacterial strains were downloaded from the Ensembl database (March 2020) (Yates, et al. 2020).

### Identification and realignment of orthologous genes

To identify the orthologs of RpoB, RpoC and the additional examined genes, we carried out BLAST (Altschul, et al. 1990) searches using the *E. coli* K12 MG1655 protein sequence as a query against the Ensmbl collection of proteomes. Best bi-directional hits were required in order to maintain an identified ortholog for further analyses. Each identified ortholog was re-aligned with the *E. coli* K12 MG1655 sequence, using the Needleman-Wunsch pairwise alignment algorithm, as implemented by the EMBOSS needle program (Rice, et al. 2000) function in Biopython (Cock, et al. 2009). This enabled us to compute the optimal alignment (including gaps) of each two sequences along their entire length. Alignments that had less than 30% overall sequence identity across their entire length were removed from consideration. To avoid biases stemming from over-representation of certain closely related groups of strains within our dataset, with identical RpoB, RpoC or other gene protein sequences, identical sequences we combined into a single representative. **Table S1** summarizes the number of alignments obtained following this procedure, for each of the studied genes.

### Z-score calculations

In order to be able to combine different genes that vary in the distributions of the conservation levels of their positions, we calculated for each gene separately a Z-score for each of its positions as:

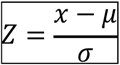

Where: x denotes the percentage of strains in which that position is conserved, μ denotes the mean percentage conservation across all positions of that protein, and σ denotes the standard deviation around that mean. Z score values were then combined across genes allowing us to compare them between positions in which adaptations occurred and all other positions.

### Mapping RpoB an RpoC positions onto the RNA polymerase molecular structure

We mapped the RpoB and RpoC positions involved in adaptation onto the three-dimensional structure of the RNA polymerase complex of *E. coli* (Protein Data Bank; 3LUO) (Opalka, et al. 2010). The mutations were mapped and visualized using PyMol (The PyMOL Molecular Graphics System, Version 2.4,

Schrödinger, LLC) and Pyrosetta (Version 2.6) (Chaudhury, et al. 2010). The distance between the residues and the active site was measured with the PyMol distancetoatom function.

In order to classify positions according to whether they fall within an annotated protein domain, the domain annotation was taken from UniPort Knowledgebase (UniProt 2021), entries P0A8V2 (RpoB) and P0A8T7 (RpoC).

## Results

### Little overlap in the RNA polymerase core enzyme positions involved in adaptation to various selective pressures

We carried out a literature survey to annotate protein positions involved in adaptation to a variety of selective pressures, within the RNA polymerase core enzyme (RNAPC) proteins RpoB and RpoC, in the model bacterium *Escherichia coli*. This resulted in the identification of 128 positions (**Table 1**). Of these positions, 38 were involved in the acquisition of antibiotic resistance. While 90 were involved in adaptation to other selective pressures, such as survival under prolonged resource exhaustion, adaptation to growth within poor nutrient media, and heat shock.

**Table 1.**
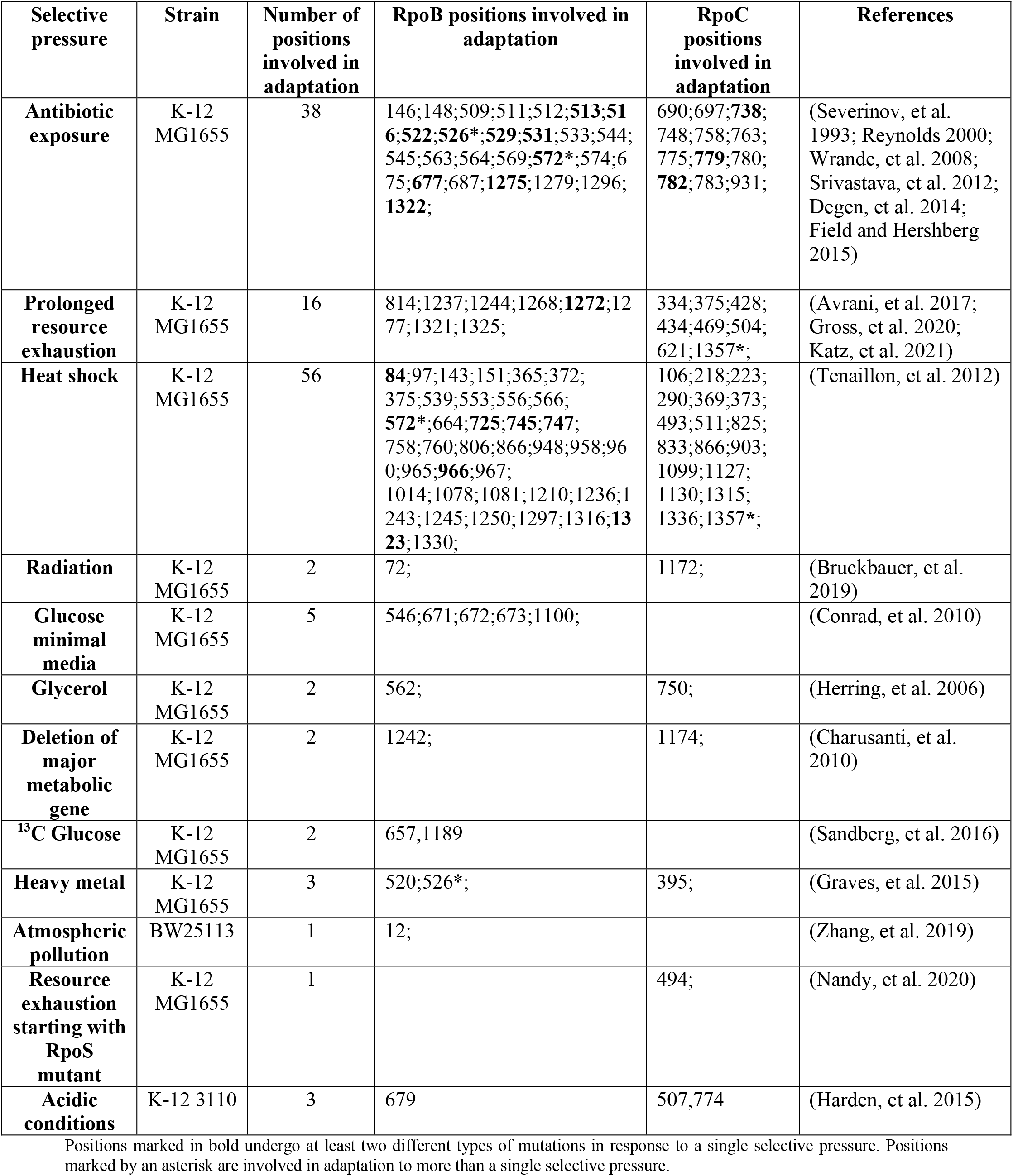
Summary of RNAPC positions involved in rapid adaptation.

It is important to distinguish between antibiotic resistance and other types of adaptation, as in the case of antibiotic resistance the reason for the occurrence of the mutations within the RNAPC is different. Specifically, antibiotic resistance adaptations occur within the RNAPC genes, because the antibiotics they confer resistance to themselves target those genes. Mutations that confer resistance are those that alter the structure of the protein so that the antibiotic can no longer effectively bind it (Spratt 1994). In contrast, an RNAPC mutation that provides an advantage under, for example, prolonged resource exhaustion likely owes its adaptive effect to the effects it has on the function of the RNAPC in regulating gene expression. Strikingly, we observe very little overlap in the RNAPC positions involved in adaptation to different selective pressures. Of the 128 positions in our dataset of positions involved in adaptation, only one is involved in adaptation to two different non-antibiotic related selective pressures, and only two are involved in both antibiotic resistance and an adaptation to a second selective pressure.

#### Adaptation tends to occur within more conserved positions of the RNA polymerase core enzyme

Next, we examined whether the 128 RpoB and RpoC positions in which adaptations were found, differed in their levels of conservation, compared to the remaining RpoB and RpoC positions. To do so, the *E. coli* K12 MG1655 RpoB and RpoC sequences were compared, using BLAST, at the protein level against a database of the full proteomes of 44,048 fully sequenced bacterial genomes. Only bidirectional best hits were maintained. Each of the identified RpoB and RpoC sequences, was then realigned at the protein level against their *E. coli* K12 MG1655 ortholog, using the Needleman-Wunsch pairwise alignment algorithm, as implemented by the EMBOSS needle program. This enabled us to compute the optimal alignment (including gaps) of each two sequences along their entire length. In order to avoid biases, resulting from over sampling of closely related bacterial strains with identical RNAPC genes, if two RpoB or two RpoC orthologs were found to be identical in their sequences, only one of the two was maintained. Finally, we filtered out alignments that had less than 30% overall sequence identity across their entire length. From the resulting 8163 RpoB alignments and 7727 RpoC alignments we calculated the percentage of strains in which each position of the *E. coli* K12 protein sequence was conserved. We than compared levels of conservation, between the 128 positions that were shown to be involved in rapid adaptation, and the remaining protein positions. This enabled us to demonstrate that positions in which adaptations are found tend to be more conserved than all remaining positions (**Figure 1**). For example, 49% and 23% of positions involved in adaptation within RpoB and RpoC respectively are conserved in 100% of the examined strains. At the same time only 19% and 13% of all other positions within these two genes are so conserved. The observed difference in levels of conservation between positions known to be involved in adaptation, and all remaining positions is statistically significant (*P* << 0.001 for RpoB, and *P* = 0.003 for RpoC, according to a one-tailed non-paired Mann-Whitney test).

**Figure 1.**
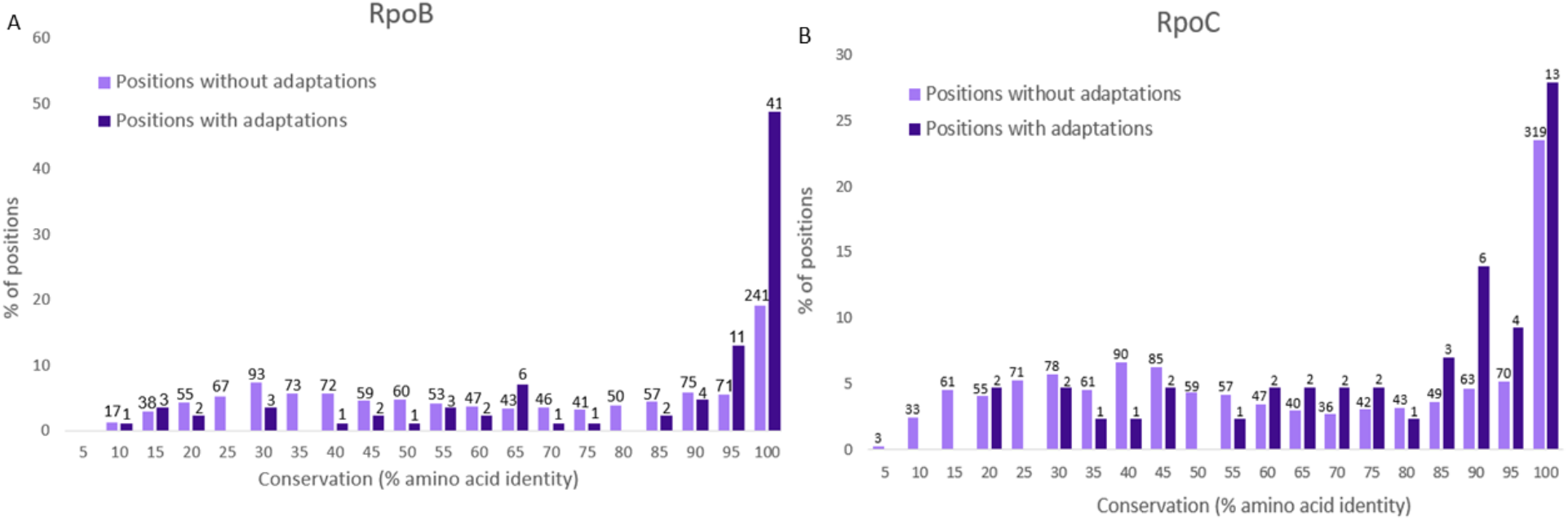
Positions in which known rapid adaptations occur tend to be more conserved than those in which no such adaptation is known. Depicted in each graph are the distributions of conservation levels of RpoB (A) and RpoC (B) positions, divided into positions in which known adaptive mutations were found to occur (dark blue) and those in which no such adaptive mutations were yet identified (lavender). Numbers above each bar indicate the numbers of positions falling within each conservation bin. Positions in which known adaptations occur are significantly more conserved for both RpoB (P << 0.001) and RpoC (*P* = 0.0064).

For adaptation to antibiotic resistance, heat shock and prolonged resource exhaustion, sufficient positions are involved to test differences in conservation separately for each selective pressure. Adaptations to prolonged resource exhaustion and adaptations leading to antibiotic resistance occur at positions that were more conserved than positions at which no adaptations were found (P << 0.001 for adaptations occurring within RpoB and *P* < 0.008, for adaptations occurring within RpoC, **Figure S1**). In contrast no significant difference was found in the conservation of positions involved in adaptation to heat shock and positions with no observed adaptation (**Figure S1**).

The *Proteobacteria* phylum to which *E. coli* belongs is one of the most well studied of bacterial phyla (Gupta 2000). As a result, sequences belonging to this phylum are likely to be over-represented within our database. As this could potentially bias results, we aimed to verify that RNAPC positions tend to be more conserved across all phylogenetic distances. To do so, we separated the 8163 of RpoB and 7727 of rpoC alignments we obtained, according to their percent identity, into 10% sized bins (e.g. 90-100%, 80-90% etc…). We then examined whether positions in which adaptations were found in *E. coli* tended to be more conserved, than positions in which no adaptations were observed, based on each group of alignments separately. In the case of RpoC, positions involved in adaptation are significantly more conserved than remaining positions, for all phylogenetic distances (P < 0.05, for all comparisons, **Figure S2**). In the case of RpoB, this was true (P < 0.001, **Figure S2**) for all but the most closely related alignments (90-100% identity, *P* = 0.1237). Our results thus demonstrate that positions involved in adaptation tend to be more conserved than other positions, across all phylogenetic distances.

#### Positions involved in adaptation tend to fall within defined functional domains

In addition to tending to be more conserved, RNAPC positions at which adaptations occur also tend to more often fall within residues belonging to defined functional domains (**Table S2**) (Ponting and Russell 2002; UniProt 2021). This is true in general, when all sites are considered together (*P* << 0.001 for RpoB and *P* = 0.0011 for RpoC, according to a Mann Whitney test), and is also true when we consider antibiotic resistance adaptations (*P* << 0.001 for RpoB and *P* = 0.0085 for RpoC) and resource exhaustion adaptations (*P* << 0.001 for RpoB and *P* = 0.0016 for RpoC) separately. However, heat shock adaptations do not show a similar tendency to be enriched within defined functional domains (*P* = 0.1405 for RpoB and *P* = 0.2645 for RpoC).

#### Positions involved in adaptation tend to be located close to the RNAPC active site

In order to further characterize the RNAPC positions involved in adaptation, we located them on the RpoB and RpoC complex solved protein structure (Protein Data Bank; 3LUO) (Opalka, et al. 2010) (**Figure 2A**). In general, positions involved in adaptation tended to be closer to the enzyme’s active site than other positions (P<<0.001, for both RpoB (**Figure 2B**) and RpoC (**Figure 2C**), and **Table S2**. see Materials and Methods). When considered separately, resource exhaustion adaptations and antibiotic resistance adaptations tend to be located closer to the complex’s active site than positions that are not involved in adaptation (*P* < 0.001, for all comparisons). At the same time, in agreement with their lower levels of conservation and lack of tendency to be enriched within functional domains, positions involved in heat shock adaptation do not display a strongly significant enrichment for proximity to the active site (*P* = 0.0253 for RpoB and *P* = 0.3509 for RpoC).

**Figure 2.**
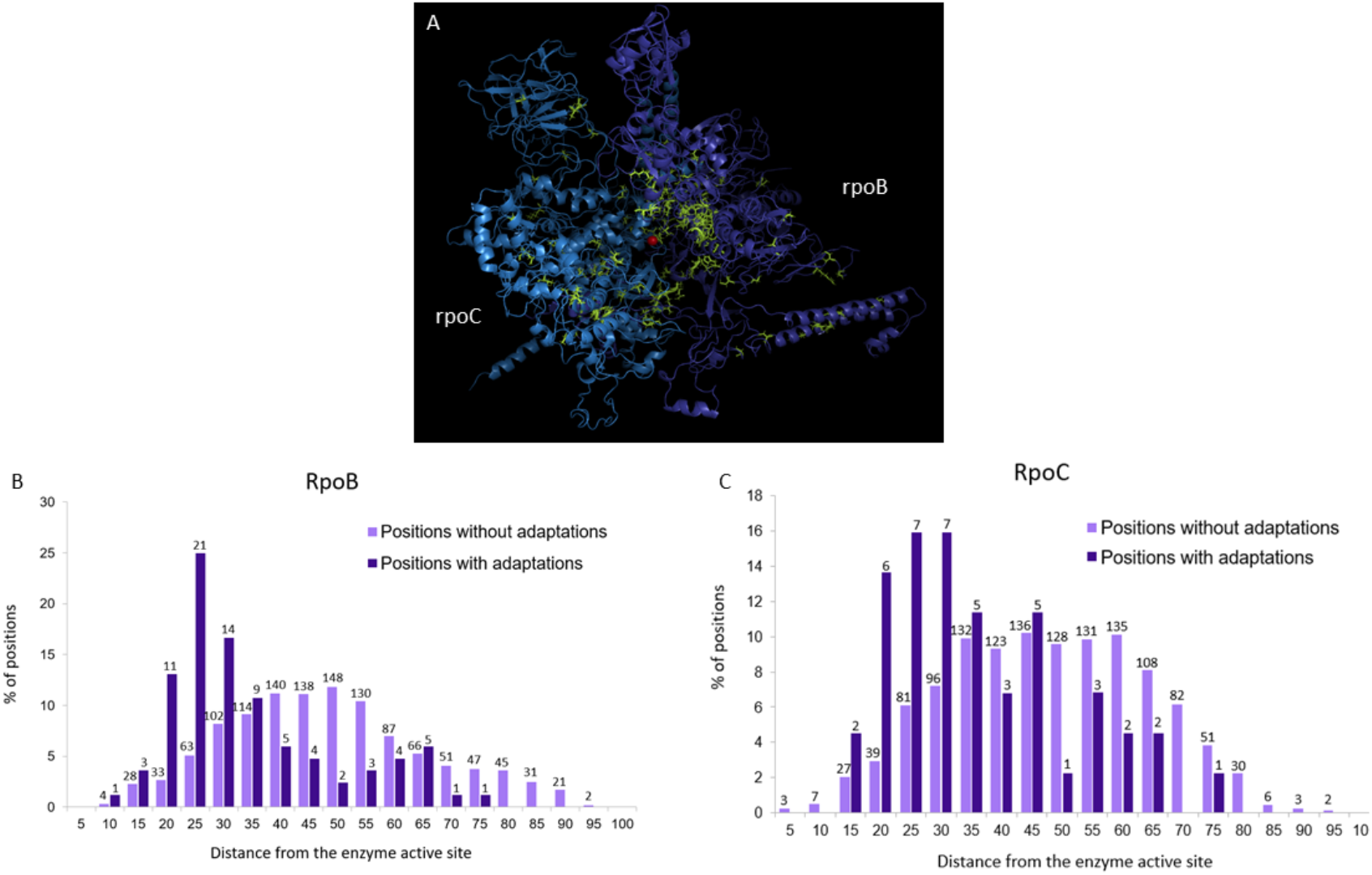
Positions involved in adaptation tend to be located closer to the RNAPC active site. (A) The solved protein structure of the RpoB-RpoC complex is presented (PDB accession 3LU0), with positions in which adaptations occur marked in green. Positions of both RpoB (A) and RpoC (B) at which known adaptations occur tend to be located significantly (P << 0.001) closer to the protein complex’s active site.

#### Excluding heatshock adaptations, positions involved in adaptation to the same condition tend to be clustered on the protein structure

Resource exhaustion adaptations (**Figure 3A**), and minimal media adaptations (**Figure 3B**) tended to each separately cluster onto distinct close regions of the protein structure. Similarly, positions involved in adaptation to acquire resistance to the same antibiotic, also tend to cluster together (**Figure 3C**). In contrast, unlike most positions involved in adaptation, positions involved in adaptation to heat shock are more dispersed over the entire RpoB and RpoC complex structure (**Figure 3D**).

**Figure 3.**
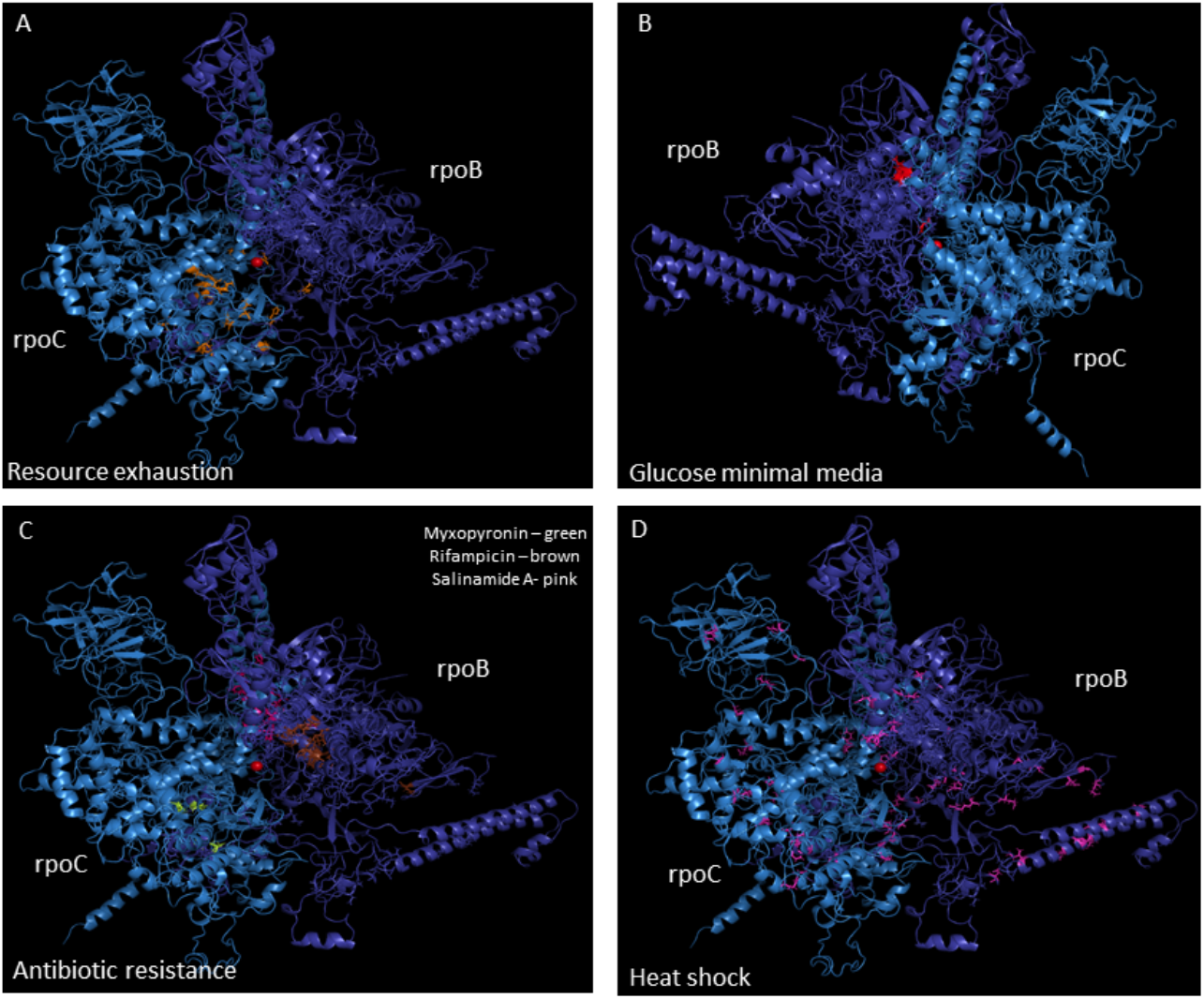
Locations of known adaptations on the solved protein structure of the RpoB-RpoC complex. The RpoB-RpoC protein structure was taken from (PDB accession 3LU0). Positions where known rapid adaptations occur marked on the structure: (A) Prolonged resource exhaustion adaptations. (B) Glucose minimal media adaptations. (C) Antibiotic resistance adaptations. (D) Heat shock adaptations.

### Resource exhaustion adaptations within additional genes, also tend to occur within more conserved positions

We have been studying *E. coli* adaptation under prolonged resource exhaustion. In our experiments we found very high levels of convergence, with mutations often occurring within the same loci, across independently evolving populations (Avrani, et al. 2017; Katz, et al. 2021). Such convergence is widely considered to be a signal of adaptation. 19 genes (other than *rpoB* and *rpoC*) were identified in which mismatch mutations occurred across all five of our populations, indicating that these mutations are adaptive under resource exhaustion **(Table S1)**. To examine whether the positions of these genes at which convergent mutations occurred tend to be more conserved than other positions of these genes, we first carried out BLAST searchers, using each gene as a query and requiring bi-directional best hits, as done before for RpoB and RpoC. Bacterial strains for which we could not find an RpoB or RpoC ortholog were removed from consideration. Alignments were then refined using the Needleman-Wunsch algorithm, with at least 30% identity required, and identical alignments were clustered together into a single alignment. The numbers of bacterial strains in which we found each gene initially, and the number of ultimate alignments we were left to work with in the end are summarized in **Table S1**.

In contrast to RpoB and RpoC, here, for each gene, only a handful of positions were predicted to be involved in adaptation. In order to obtain sufficient power to examine whether positions likely involved in adaptation tended to be more conserved, we had to therefore combine data across our 15 genes. To do so, while not biasing our results due to differences in overall conservation between the proteins, we normalized within each gene the calculated levels of conservation of each position by calculating a Z-score (**Table S3, Materials and Methods**). Once conservation levels were normalized, they could then be combined across genes. We found that the positions within convergently mutated genes, in which resource exhaustion mutations occurred were significantly more conserved than remaining positions within the same genes (*P* << 0.001, according to a one-tailed non-paired Mann -Whitney test).

## Discussion

We demonstrate that adaptation within the RNAPC and additional housekeeping genes tends to occur within the most conserved and evolutionarily constrained positions of these highly conserved proteins. This raises a conundrum: How is it possible for these positions to change so rapidly in response to a variety of selective pressures, yet remain so highly conserved over longer evolutionary time scales? The answer to this question may relate to pleiotropy (Cooper and Lenski 2000; MacLean, et al. 2004; Kvitek and Sherlock 2013). The specific changes to the RNAPC and additional master regulatory genes that are adaptive under a specific condition, may have strongly deleterious pleiotropic effects under many, if not all other conditions. After all, master regulators in general, and the RNAPC in particular regulate the expression of large chunks of the transcriptome. In lab experiments, bacteria are generally exposed to relatively simple, strong and constant selective pressures. The selective pressures faced within more natural environments are likely far more complex, with several different factors exerting contradictory pressures simultaneously and / or with selective pressures that change with time. Adaptations of the kind that arise so easily during lab evolution, may not be so easily permitted within natural environments, due to their pleiotropic effects. Additionally, if such adaptations do occur in response to a specific set of conditions, they may prove to be highly transient, rapidly decreasing in frequency once conditions change. Supporting this, we recently demonstrated such transience of RNAPC adaptations arising under resource exhaustion. We showed that bacteria exposed to prolonged resource exhaustion adapt via mutations to the RNAPC, but that these mutations do not tend to fix across an entire population (Avrani, et al. 2020). Since these mutations carry pronounced costs to fitness under conditions of resource abundance and rapid growth, once bacteria are transferred into fresh media, rare clones that do not carry an RNAPC adaptation outcompete the clones that do carry these adaptations, leading to rapid reductions in the frequencies the RNAPC adaptations (Avrani, et al. 2020). If RNAPC adaptations are in general highly transient, it may explain why the sites in which they occur can remain largely conserved, when one examines longer evolutionary timescales.

We find no overlap in the positions of the RNAPC involved in adaptation to various selective pressures, indicating that different selective pressures demand different specific changes to the RNAPC. We further find that for most conditions, sites involved in adaptation tend to be clustered on the protein structure. Combined, these results strongly suggest that for most conditions, very specific changes to the RNAPC are adaptive.

The specificity of sites involved in adaptation under each selective pressure may suggest that adaptation to each pressure occurs through changing a specific function of the RNAPC, different from the function changed in response to a different selective pressure. In other words it is possible that the reason that specific positions are involved in adaptation to condition A, while different ones are involved in adaptation to position B, is that the condition A adaptations drive specific gene expression changes adaptive under condition A, while position B adaptations drive different changes to gene expression, adaptive under condition B. A second possible reason for the specificity of the adaptive mutations, is that while the same ultimate adaptive outcome is always reached, the way to reach that outcome is condition dependent. In other words, the adaptive outcome may involve the same specific change to transcriptional kinetics, under both conditions A and B. However, due to changes in the structure of the transcriptional regulatory network under the different conditions, inducing the adaptive end result might involve different mutations in condition A, compared to condition B. Finally, high levels of specificity may also be explained not through the need to change a specific function, but through the need to prevent antagonistically pleiotropic effects (Cooper and Lenski 2000; MacLean, et al. 2004; Kvitek and Sherlock 2013). It is possible that under different selective pressures, adaptive mutations always carry out a similar function (e.g. altering the kinetics of transcription in a similar manner), yet different specific mutations can be tolerated under each condition, due to differences in their pleiotropic effects. In other words, it is possible that the positions involved in adaptation to condition A are ones in which mutations can alter the kinetics of transcription, in a manner that is adaptive across conditions, while at the same time minimizing changes to gene expression that would be specifically harmful under condition A. A combination of all these explanations may also be possible.

Some clues as to the consequences of the RNAPC adaptive mutations may be gleaned from their location on its structure. In the case of antibiotic resistance adaptations, the reasons for their location are easy to predict: Resistance adaptations will likely occur within the same region of the complex that the antibiotic binds, and work by reducing the ability of the antibiotic to bind its target. When it comes to other types of adaptations it is less straight forward to predict their precise adaptive effect. Conard et al. revealed the adaptive role of minimal media RNAPC adaptations in altering transcriptional kinetics, by decreasing the longevity of open complex. Here, we found that resource exhaustion RNAPC adaptations are located within the complex’s clamp domain, which has been implicated in involvement in several crucial aspects of the transcription process, including transcription initiation and elongation (Duchi, et al. 2018).

Mutations to the RNAPC involved in adaptation to heat shock, behave differently than those involved in adaptation to other conditions. While adaptation to heat shock appears to occur within the RNAPC in a highly convergent manner, many different specific mutations were found to occur across independently evolving populations (Tenaillon, et al. 2012). This stands in contrast adaptation to prolonged resource exhaustion, where the same specific sites tend to be mutated across independently evolving populations (Avrani, et al. 2017; Katz, et al. 2021). The many heat shock RNAPC adaptations do not tend to occur within significantly more conserved positions, are not enriched within known functional domains or in proximity to the active site, and they are not clustered together on the complex’s structure. This appears to suggest that unlike adaptations to other conditions, heat shock adaptations may be acting through a less specific mechanism, affecting some more general trait of the RNAPC, that does not require changes to very specific sites, located at the heart of the complex. Intriguingly, studies into the effects of RNAPC heat shock adaptations suggested that these adaptations change the expression of hundreds of genes back towards the transcriptional program of pre-stressed bacteria (Rodriguez-Verdugo, et al. 2016). It will be interesting to understand how this can be achieved by such a large variety of non-specific mutations, located all over the RNAPC’s structure.

## Acknowledgments

This work was supported by an ISF grant (No. 1860/21, to R.H.) and by the Rappaport Family Institute for Research in the Medical Sciences (to R.H.). The described work was carried out in the Rachel & Menachem Mendelovitch Evolutionary Process of Mutation & Natural Selection Research Laboratory.

