## Supplementary material for "Rapid adaptation often occurs through mutations to the most highly conserved positions of the RNA polymerase core enzyme": Table S1

**Table S1**: Number of alignments left after filtration

| **Gene** | **Number of alignments Prior to filtration** | **Number of alignments following filtration** |
| --- | --- | --- |
| ***asmA*** | 16,748 | 966 |
| ***crp*** | 31,745 | 781 |
| ***cytR*** | 28,755 | 820 |
| ***dcuA*** | 43,985 | 1652 |
| ***deoR*** | 14,485 | 730 |
| ***dppA*** | 17,556 | 2,175 |
| ***fadR*** | 11,916 | 648 |
| ***glpF*** | 34,254 | 2380 |
| ***gltS*** | 21,596 | 1,707 |
| ***kgtP*** | 20,173 | 2,407 |
| ***oppA*** | 21,726 | 1,993 |
| ***paaX*** | 17,384 | 720 |
| ***prc*** | 34,870 | 2516 |
| ***putP*** | 23,971 | 3,148 |
| ***rimJ*** | 28,682 | 1565 |
| ***rpoA*** | 43,375 | 4,457 |
| ***rpoD*** | 42,687 | 5,468 |
| ***sstT*** | 7,992 | 977 |
| ***sucC*** | 32,306 | 4,722 |
