## Supplementary figures and images for "Rapid adaptation often occurs through mutations to the most highly conserved positions of the RNA polymerase core enzyme"

### Figure S1

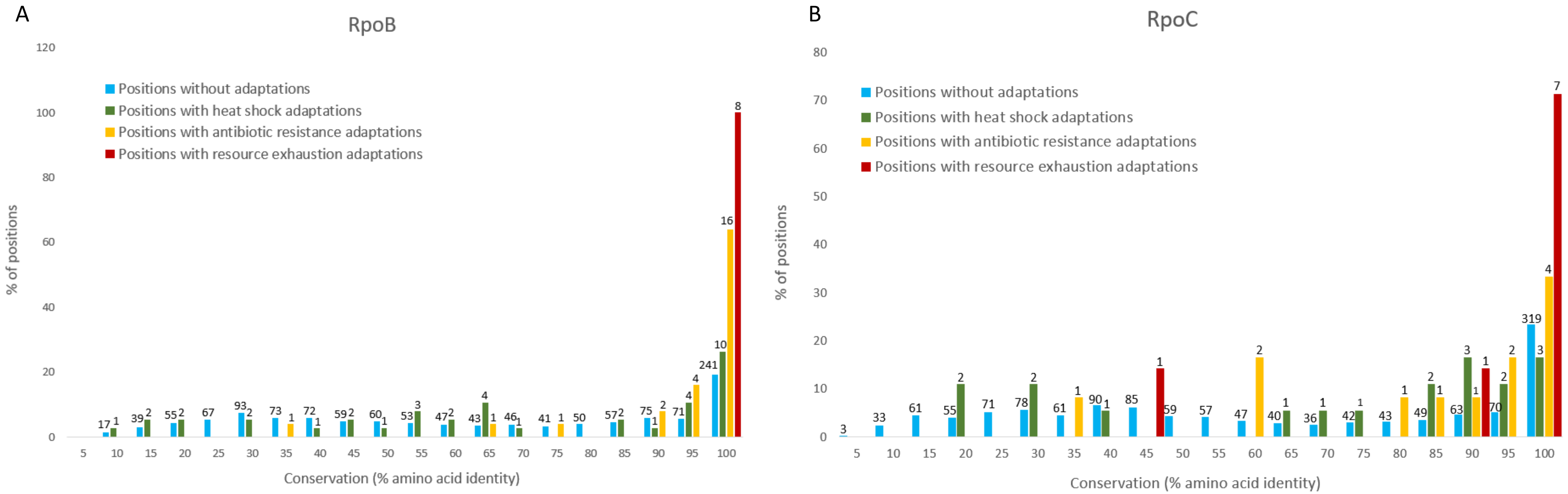

### Figure S2

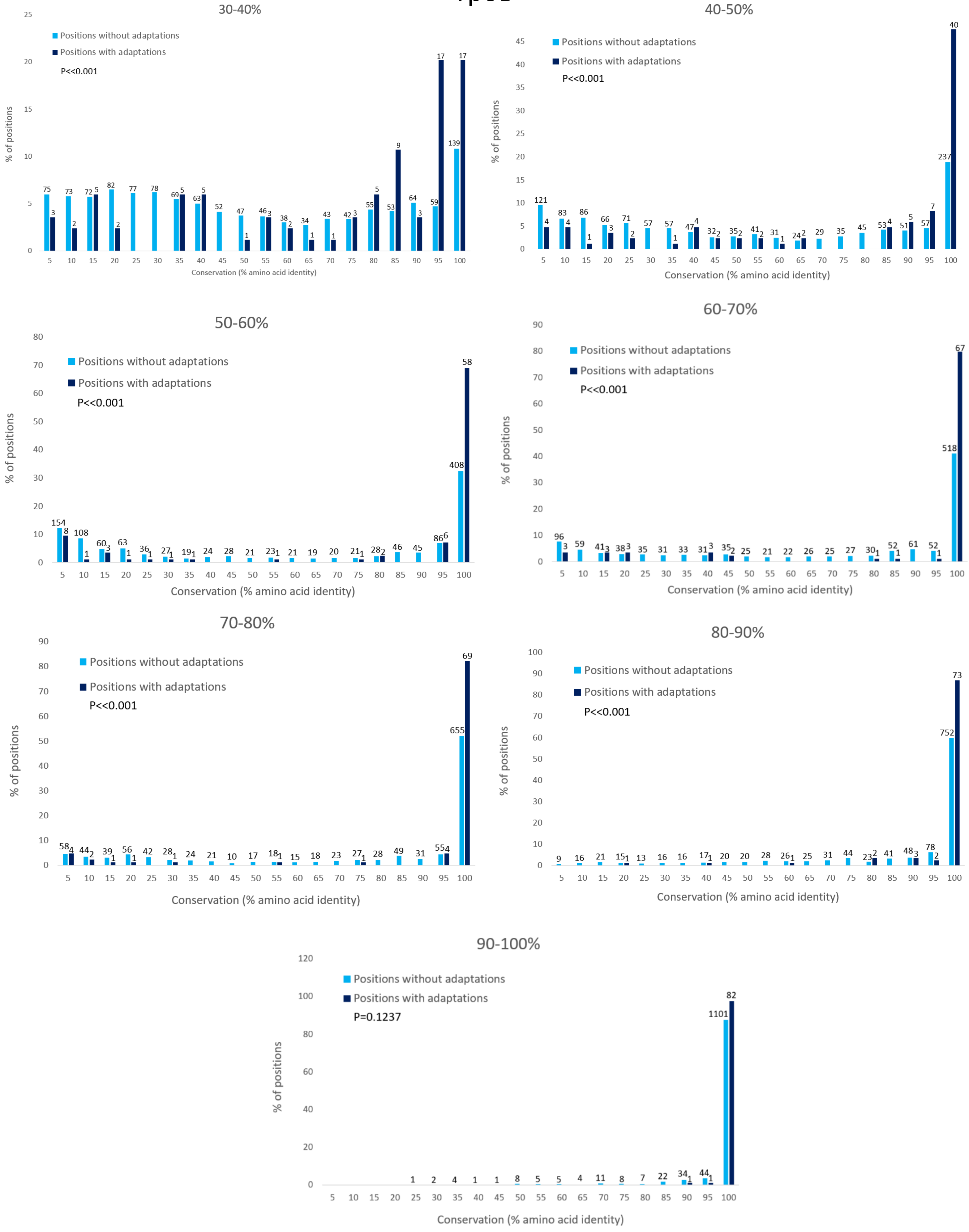

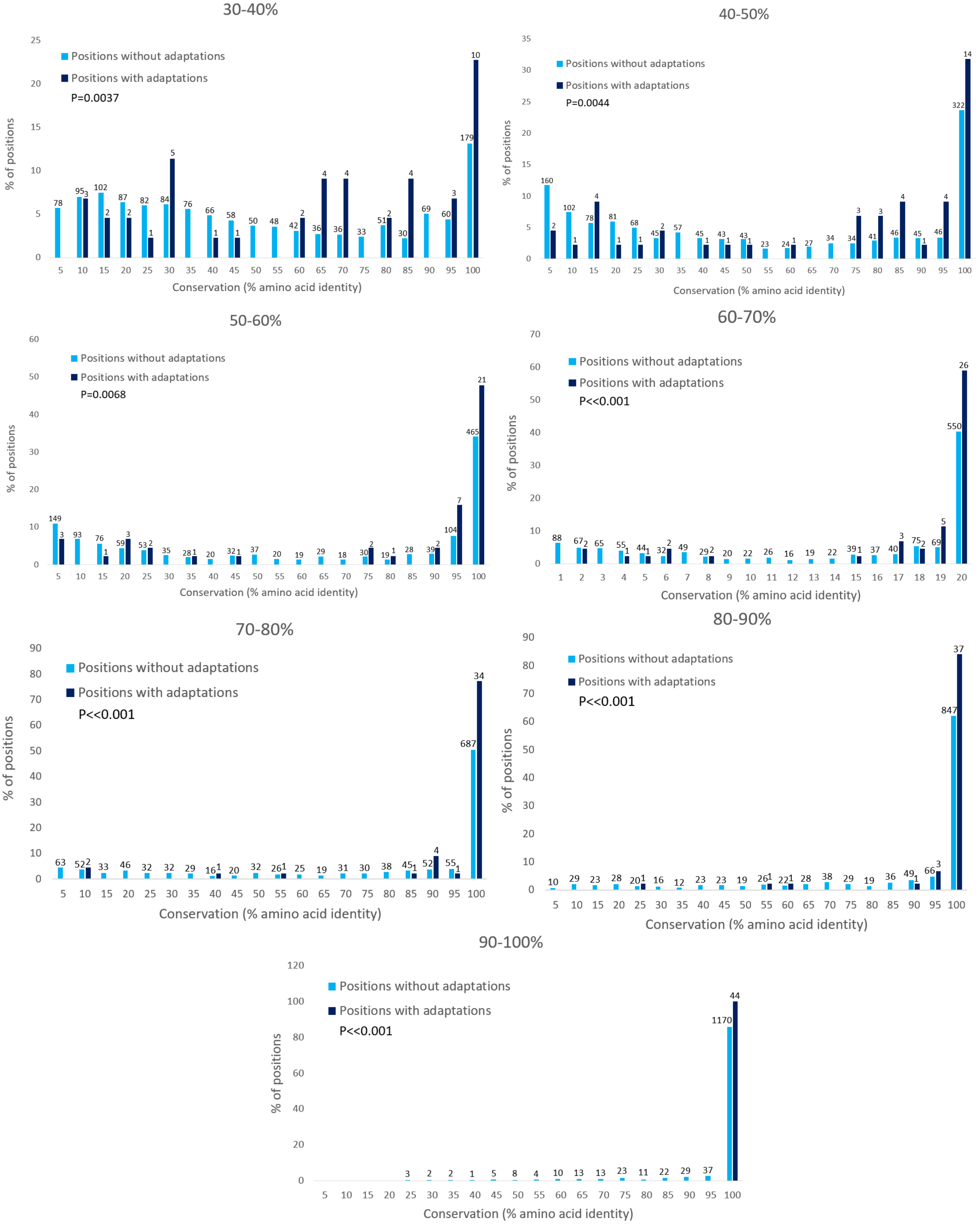
